# Evolutionary rates in human amyloid proteins reveal their intrinsic metastability

**DOI:** 10.1101/2022.09.07.506994

**Authors:** Diego Javier Zea, Juan Mac Donagh, Guillermo Benitez, Cristian Guisande Donadio, Julia Marchetti, Nicolas Palopoli, María Silvina Fornasari, Gustavo Parisi

## Abstract

The emerging picture of protein nature reveals its intrinsic metastability. According to this idea, although a protein is kinetically trapped in a local free energy minimum that defines its native state, those kinetic barriers can be overcome by a complex mixture of the protein’s intrinsic properties and environmental conditions, promoting access to more stable states such as the amyloid fibril. Proteins that are strongly driven towards aggregation in the form of these fibrils are called amyloidogenic. In this work we study the evolutionary rates of 81 human proteins for which an *in vivo* amyloid state is supported by experiment-based evidence. We found that these proteins evolve faster when compared with a large dataset of ∼16,000 reference proteins from the human proteome. However, their evolutionary rates were indistinguishable from those of secreted proteins that are already known to evolve fast. After analyzing different parameters that correlate with evolutionary rates, we found that the evolutionary rates of amyloidogenic proteins could be modulated by factors associated with metastable transitions such as supersaturation and conformational diversity. Our results showcase the importance of protein metastability in evolutionary studies.

## Main Manuscript

Toxicity derived from protein misfolding and aggregation has been the core concept to explain the strong anticorrelation between expression level and evolutionary rate observed in most organisms studied so far (Pál et al. 2001; Drummond and Wilke 2008). According to the translational robustness hypothesis (Drummond et al. 2005) and more recent extensions such as the misfolding avoidance hypothesis (Yang et al. 2010), highly expressed proteins evolve slowly due to an increased selection pressure against misfolding or aggregation. As a derivation of these hypotheses, highly expressed proteins are expected to be thermodynamically more stable (Drummond and Wilke 2008). However, recent studies have not found any correlation between experimentally measured protein stability and evolutionary rates (Plata and Vitkup 2018; Usmanova et al. 2021). Therefore, the proposed causal relationship between aggregation and slow evolutionary rates is still controversial.

Amyloidogenic proteins tend to be soluble and globular but undergo substantial conformational changes to aggregate as amyloid fibrils under certain circumstances. They don’t show evident sequence, fold, or function similarities, but their aggregation propensity has been correlated with different sequence determinants such as hydrophobicity, β-strand propensity, and net charge (López de la Paz and Serrano 2004). Accumulation of residues with these properties into one or several short aggregation-prone regions (APR) was extensively used to predict amyloid aggregation propensity at the proteomic level (Fernandez-Escamilla et al. 2004; Ventura et al. 2004; Maurer-Stroh et al. 2010; Walsh et al. 2014; Zambrano et al. 2015). Different processes such as mutations, physiological stress, and stability loss expose these APRs to the solvent, increasing the chances that they reach a stable state by self-assembling into amyloid fibrils (Beerten et al. 2012; Buck et al. 2013).

Amyloids have long been associated with the occurrence of over 50 diseases in humans by aggregation of proteins like transthyretin in the transthyretin amyloidosis (da Costa et al. 2015), amyloid precursor and tau proteins in Alzheimer’s disease (Murrell et al. 2000), alpha-synuclein in Parkinson’s disease (Stefanis 2012), and prion protein in the spongiform encephalopathies (Weissmann et al. 2002). Considering the extreme conformational changes in which most amyloidogenic proteins are involved, we hypothesized that they would show a particular substitution pattern during evolution due to a high selective pressure (Zea et al. 2013). To explore this idea, we assembled two large datasets of paired orthologous proteins between human and mouse (*Homo sapiens - Mus musculus*, with 16,589 proteins) and between human and cattle (*Homo sapiens - Bos taurus*, with 15,974 proteins). We also collected 11,789 common orthologous proteins in seven species related by a single phylogenetic tree *(Homo sapiens;* its close relatives *Pan troglodytes, Macaca fascicularis* and *Rhinopithecus bieti;* and three other mammals, *Mus musculus, Bos taurus*, and *Sus scrofa)*. All these orthologs were extracted from the OMA database (Altenhoff et al. 2018, see Supplementary Information for further details). Using the Amypro database of validated amyloid precursor proteins (Varadi et al. 2018) and manual curation from recent bibliography (see Supplementary Information), we identified 81 human proteins with experimental evidence of aggregation into an amyloid form. These were defined as the “amyloids” dataset, in opposition with the rest of the proteins without evidence of amyloid fibril formation *in vivo*, which we took as the “reference” dataset. Evolutionary rates, expressed as the number of non-synonymous substitution rates per non-synonymous site (*dN*), were estimated for pairs of homologous proteins using the program YN00 from the package PAML (Yang 1997), additional information in Supplementary files). In the case of the seven species dataset, the program CODEML was used to estimate *dN*, using the same reference tree.

Using the dataset of human - mouse orthologs, we found a higher *dN* for the amyloid proteins (Wilcoxon rank-sum test p-value < 0.01) in comparison with the reference dataset (Figure 1) (All results shown here are for the human - mouse dataset, which are representative of the results obtained for the others; see Supplementary Figure 1 for details). Unexpectedly for such fast-evolving proteins, amyloids also show higher expression levels compared with the reference proteins (Wilcoxon rank-sum test p-value < 0.05), as well as higher abundance (Wilcoxon rank-sum test p-value < 0.001, see Figure 1). These findings do not agree with the well-established observation that highly expressed proteins evolve slowly (Drummond et al. 2005; Drummond et al. 2006; Park and Choi 2009; Zhang and Yang 2015). As extracellular location (Feyertag et al. 2017) and the presence of disulphide bonds (Feyertag and Alvarez-Ponce 2017) have been found to characterize fast-evolving proteins, we split the dataset according to these variables. We annotated proteins as having disulphide bonds following UniProt annotations, and we used the database MetazSecKB (Meinken et al. 2015) to define subsets of membrane-related, intra- and extra-cellular proteins. We found that amyloidogenic proteins evolve as fast as secreted proteins (with or without disulphide bonds) (Wilcoxon rank-sum test P > 0.01). Also, we did not observe any influence of disulphide bonds in the evolutionary rates of amyloid datasets (Figure 2).

**Figure 1:**
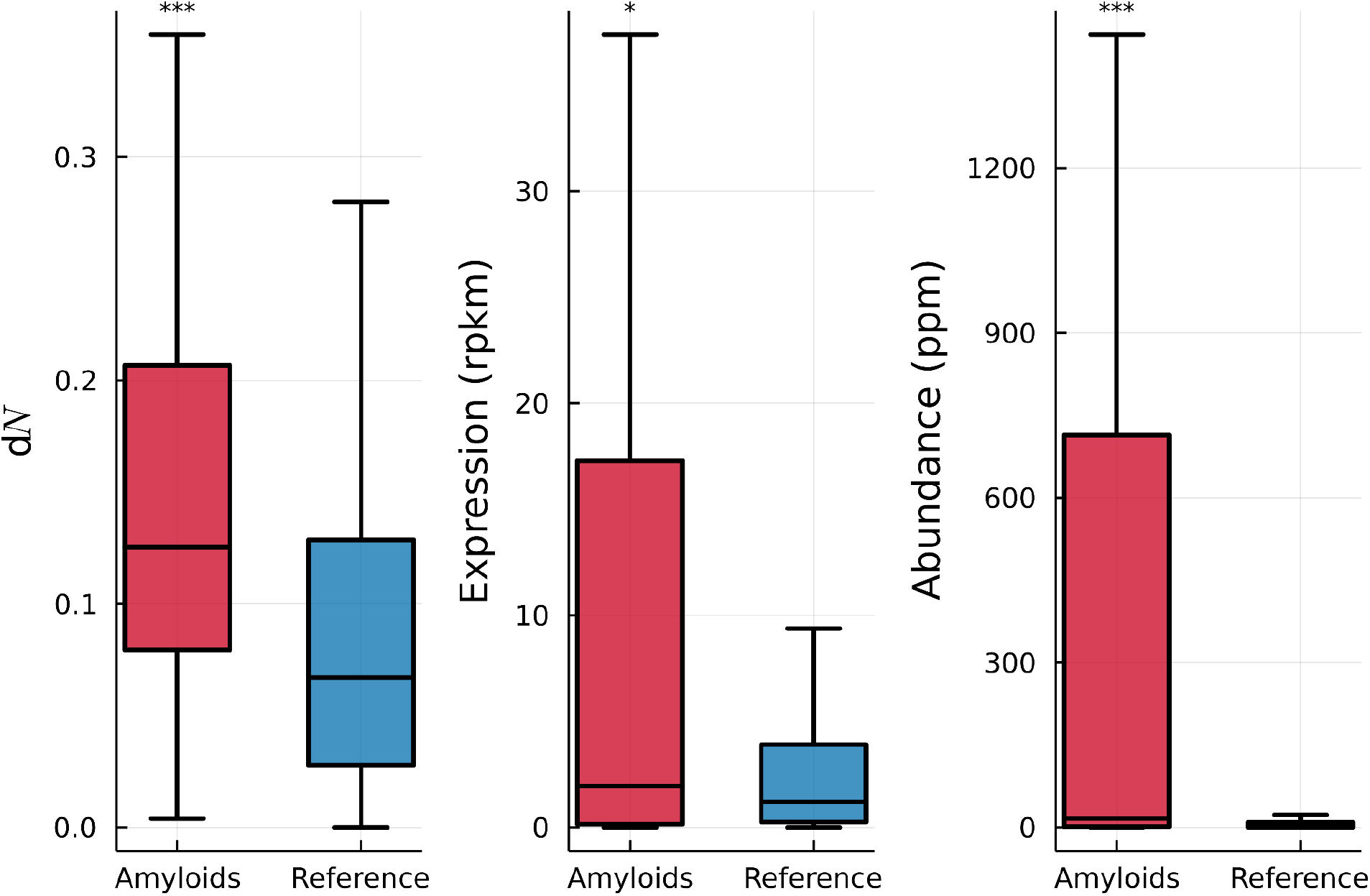
Amyloidogenic proteins evolve faster than the rest of the proteins (left panel, Amyloids n = 62, Reference n = 14,502), are more expressed (center panel, Amyloids n = 72, Reference n = 17,087), and more abundant (right panel, Amyloids n = 77, Reference n = 17,345). Stars indicate Wilcoxon rank-sum test results: ***: p-value<= 0.001; ** p-value <= 0.01; * p-value <= 0.05; ns: p > 0.05.

**Figure 2:**
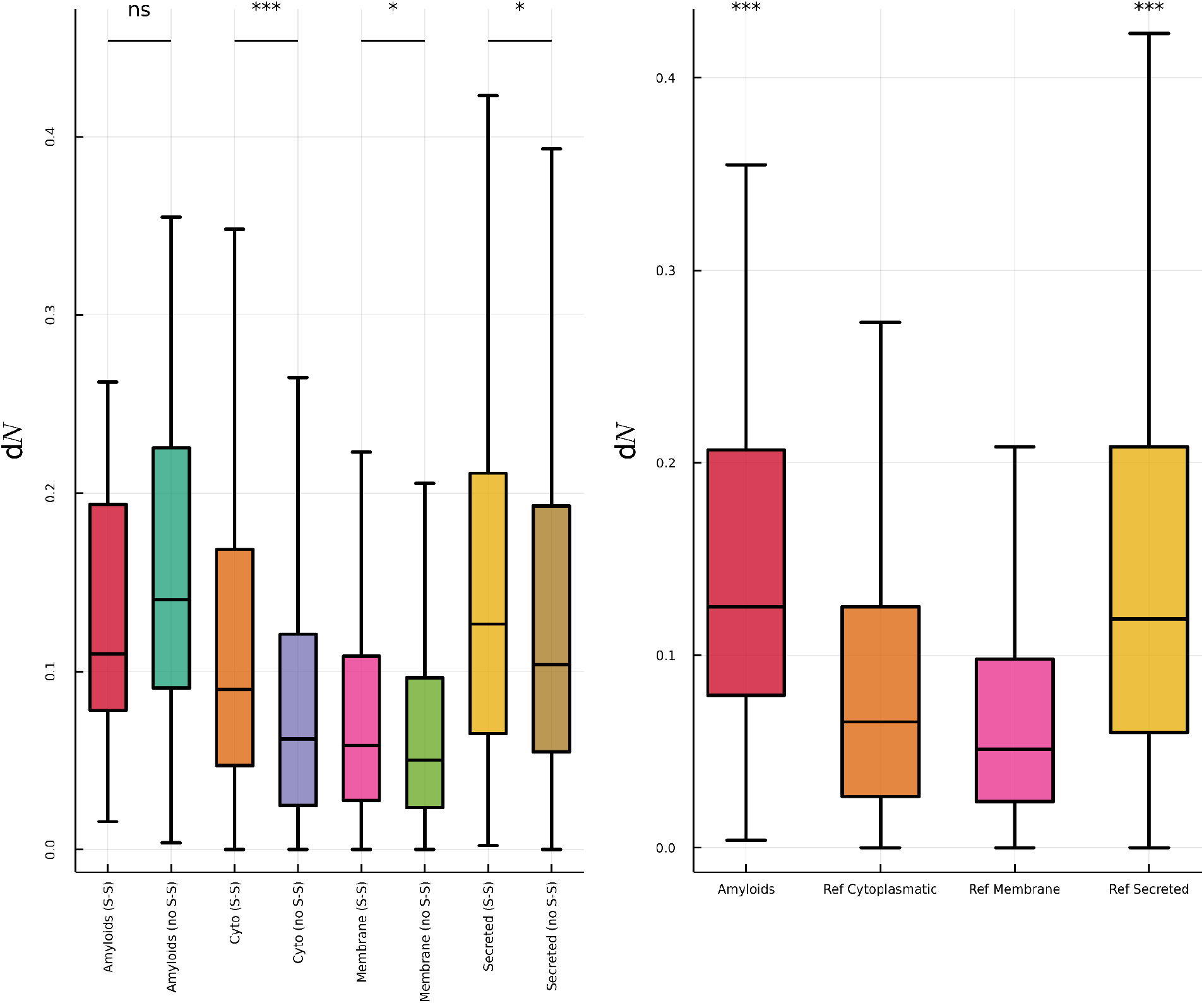
Disulfide bonds (S-S) do not relatively affect the evolutionary rates of amyloid proteins, therefore, these datasets are indistinguishable from each other, but they affect the rates of the rest of our reference dataset (left panel: Amyloids with S-S = 30, Amyloids no S-S = 27, Cytoplasmic with S-S = 1303, Cytoplasmic no S-S = 9751, Membrane with S-S = 274, Membrane no S-S =1468, Secreted with S-S = 860, Secreted no S-S = 380). However, amyloids and secreted proteins in general evolve faster than membrane and cytoplasmic proteins, but they show no statistical difference between them (right panel, both populations are faster when compared with cytoplasmic and membrane proteins). Stars indicate Wilcoxon rank-sum test results: ***: p-value<= 0.001; ** p-value <= 0.01; * p-value <= 0.05; ns: p > 0.05. “Ref” stands for the Reference dataset. “S-S” and “no S-S” indicate proteins with and without disulfide bonds, respectively.

Secreted proteins are synthesized into the endoplasmic reticulum. In the secretion process, they interact with chaperones and folding assistant enzymes to promote the right folding process and recognize misfolded proteins (Braakman and Hebert 2013). These quality controls have been pointed out as responsible for the lack of correlation between expression level (or protein abundance) and evolutionary rates, likely due to a relaxed pressure in translation errors (Feyertag et al. 2017). The same control mechanisms could be responsible for the fast evolutionary rates of secreted proteins (Figure 2) due to the buffering effects of slightly destabilizing substitutions imposed by chaperones on their clients (Tokuriki and Tawfik 2009; Williams and Fares 2010; Alvarez-Ponce et al. 2019). As about 70% of the amyloidogenic proteins in our dataset are secreted proteins, we assumed that the same mechanisms could also explain their fast evolutionary rates. It is well documented that similar quality control mechanisms are applied to cytoplasmic amyloids; for example, chaperone systems are activated to avoid amyloid fibril toxicity, avoiding and, in some cases, reverting the fibril formation (Yerbury et al. 2007; Wilson et al. 2008; Månsson et al. 2014; Landreh et al. 2015; Wentink et al. 2019; Hervás and Oroz 2020). Accordingly, we did not find a statistically significant difference between the evolutionary rates of secreted and cytoplasmic amyloidogenic proteins (Wilcoxon rank-sum test, p-value = 0.8061, see Supplementary Figure 2).

Why do amyloidogenic proteins aggregate while secreted proteins don’t, considering that both are under a strong fold quality control? As protein stability has been used to characterize aggregation and misfolding (Knowles et al. 2014), we mapped both sets of proteins in the ProTherm database (Kumar et al. 2006). Using Gibbs unfolding free energies (ΔG^0^) and melting temperatures (T_m_) as a measure of protein stability (Becher et al. 2018) we did not find any significant difference between amyloidogenic and secreted proteins (see Supplementary Figure 3). Also, as derived from their APR occurrence (estimated from TANGO predictions) (Fernandez-Escamilla et al. 2004), we found counterintuitive results because secreted proteins show the same APR content than the amyloid dataset (see Supplementary Figure 3).

Although Gibbs unfolding free energies and melting temperatures have been extensively used to estimate protein stability and the tendency to unfold and aggregate, measuring such tendencies is more complex due to the metastable nature of proteins (Baldwin et al. 2011). Metastability is a condition characterizing a protein state that is kinetically stable but has a thermodynamic tendency to change. The native state of proteins is described by a set of structures that are kinetically trapped in a local free energy minimum under physiological conditions. However, metastability means that the kinetic barriers can be overcome under certain cellular conditions that allow the native state to adopt more stable arrangements, such as the amyloid fibrils (Baldwin et al. 2011) or kinetically controlled conformations associated with protein function (Lee et al. 2000; Ghosh and Ranjan 2020). As Gibbs energy landscapes strongly depend on protein concentration, we can use protein abundance and expression level as proxies for protein aggregation (Ciryam et al. 2015; Buell 2022). Also, large conformational changes and flexibility, cavities with unusual inter-residues interactions, post-translational modifications, and protein cleavage have been associated with transitions to metastables states (Ghosh and Ranjan 2020). High protein concentration can also increase the absolute concentration of low populated conformations, perhaps partially unfolded and exposing APR regions, that could trigger fibril formation (Faravelli et al. 2022). Taking expression level, disorder (or protein flexibility), protein conformational diversity and protein abundance as proxies for the aggregation propensity, we found that those parameters are significatively higher for the amyloid dataset than for the secreted proteins (Figure 3). As in previous works (Zea et al. 2013), we measured the conformational diversity of a protein using the normalized maximum RMSD (max RMSD100,Carugo and Pongor 2001) calculated between all pairs of its known structural conformers obtained from the CoDNaS database (Monzon et al. 2016). When partial correlations were estimated to control the effect on *dN* over different confounding parameters in the amyloid dataset (n = 25, see Supplementary Table 1), using log transformed values we found significant correlations of *dN* with max RMSD100 (Pearson’s ρ = 0.49, p-value < 0.05), abundance (Pearson’s ρ = 0.47, p-value < 0.05) and protein-protein interactions (Pearson’s ρ = -0.495, p-value < 0.05).

**Figure 3:**
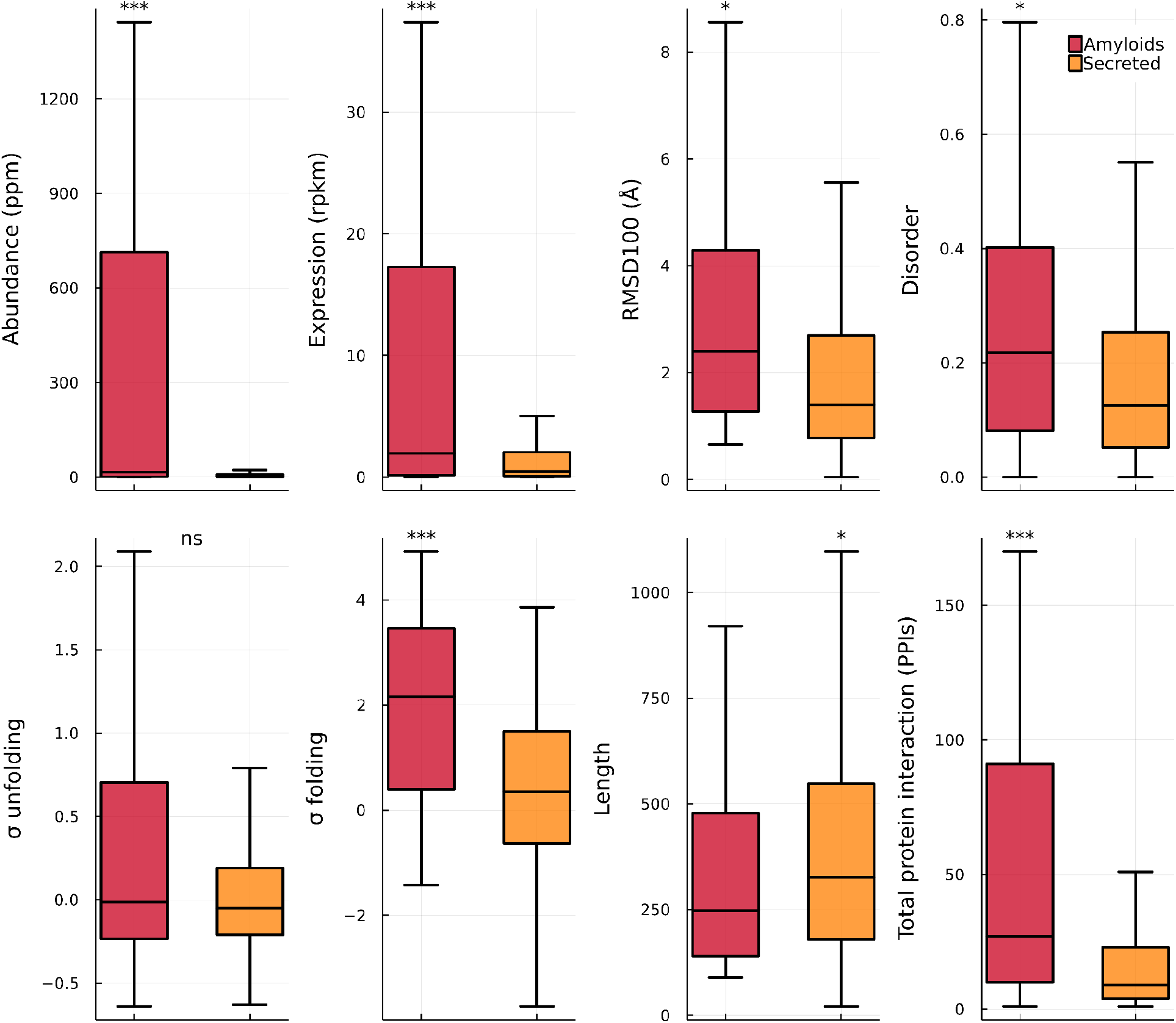
Observed differences between amyloids and secreted proteins (Amyloids = 77, Secreted = 1829, except for RMSD100, Amyloids = 34, Secreted = 389). Even though they had similar evolutionary rates, amyloids behave differently than secreted proteins with regards to variables that are correlated to evolutionary rates. Stars indicate Wilcoxon rank-sum test results: ***: p-value<= 0.001; ** p-value <= 0.01; * p-value <= 0.05; ns: p > 0.05

When protein abundance and max RMSD100 were combined in a linear regression model using the log transformations of the variables, we found that they explain ∼24% of the *dN* variance (p-value <0.02, see Supplementary Table 2). The observed variance on *dN* increased to ∼26% by including protein-protein interactions in the model (p-value = 0.073).

As explained above, conformational diversity and protein abundance can be taken as a proxy to metastability transitions. We also analyzed the influence of supersaturation, a more specific parameter associated with transitions in metastable proteins. The supersaturation score (σ_f_ and σ_u_ for the folded and unfolded states of proteins, respectively) is a linear combination of aggregation tendency and protein concentration. It identifies proteins with a high tendency to aggregate and was successfully used to mark pathological aggregation (Ciryam et al. 2013). Proteins are said to be supersaturated when their concentrations exceed their solubility levels, increasing the propensity to aggregate. We found that σ_f_, but not σ_u_, is higher in the amyloid dataset when compared with secreted proteins. We also observed a high correlation between *dN* and σ_f_ (Spearman’s ρ=0.639, p-value < 0.01, n= 17, power = 0.81). Although the number of amyloidogenic proteins is reduced due to the availability of σ_f_, when we added σ_f_ to the previous model we found that the explained variance on *dN* was ∼38% (Adjusted R^2^ = 0.388, p-value = 0.012, see Supplementary Table 3). Applying the same model to secreted proteins gave a non-significant linear model (Adjusted R^2^ = 0.008, p-value = 0.198).

To summarize, we have explored the evolutionary rates of a population of human proteins (n=81) with experimentally based evidence of aggregation as amyloid fibrils. We found that amyloidogenic proteins evolve faster than the reference dataset, in spite of being highly expressed and abundant. Although we did not observe significant differences in the evolutionary rates of secreted and amyloidogenic proteins (Figure 2), we found substantial differences in other features. Firstly, conformational diversity is higher in amyloidogenic proteins, also evidenced by the higher presence of disordered or highly flexible regions (Figure 3). This increased conformational diversity could raise the chances of exposing APRs in slightly unfolded conformers, driving the protein towards fibril formation. Secondly, amyloidogenic proteins are more abundant and expressed than secreted proteins (Figure 3). Protein concentration and solubility and their effects on driving amyloid formation were extensively studied before (see, for example, Ciryam et al. 2013). To remain soluble, abundant proteins require the constant support of quality control mechanisms such as molecular chaperones. This assistance and the well-documented participation of chaperones to avoid and revert amyloid formation could explain the accelerated evolutionary rates in amyloidogenic proteins allowed by the buffering effects of mild destabilizing substitutions (Tokuriki and Tawfik 2009). Furthermore, the positive correlation between max RMSD100 and evolutionary rates indicates that an increase in conformational diversity speeds up evolutionary rates, compatible with a higher propensity to aggregate. Thirdly, a linear model combining an intrinsic protein parameter like a measure of the conformational diversity (e.g., max RMSD100) with a cellular condition (like the supersaturation score) better explains the variation in the evolutionary rates observed in amyloidogenic proteins but not in secreted ones (Supplementary Table 3).

Emerging evidence suggests that amyloidogenic proteins represent a metastable subproteome (Olzscha et al. 2011; Ciryam et al. 2013; Kundra et al. 2017), strongly driven towards the formation of amyloid fibrils. In this particular set of proteins, intrinsic properties along with cellular conditions can release the kinetic trapping of their native states towards more stable forms such as amyloid fibrils. Our results show that evolutionary rates reflect this particular behavior, showcasing the importance of metastability above other modulating factors. In the future, it could be interesting to evaluate protein metastability as a general modulating factor of evolutionary rates at the proteome level.

## Funding / Acknowledgements

MSF, NP and GP are researchers and GIB, JM, JMD and CEGD are PhD fellows from Consejo Nacional de Investigaciones Científicas y Técnicas (CONICET). DJZ is funded by PRAIRIE (N° LS: 233070). This work was supported by Universidad Nacional de Quilmes (PUNQ 1004/11 and PP 2019), Agencia Nacional de Promoción Científica y Tecnológica (PICT-2014-3430) and the European Union’s Horizon 2020 Research and Innovation Programme (Grant Agreement N° 778247 and N° 823886). The funders had no role in study design, data collection and analysis, decision to publish, or preparation of the manuscript. The authors would like to thank Lic. Nahuel Escobedo for his help with data acquisition and Dr. Laura Acion for her feedback on the statistical analysis.

## Notes

### Competing Interest Statement

The authors have declared no competing interest.

