## Supplementary information for "Evolutionary rates in human amyloid proteins reveal their intrinsic metastability"

### Obtaining the dataset

We used the OMA database (Altenhoff et al. 2018) to obtain three datasets containing pairs of orthologs between human proteins and mouse (*Homo sapiens-Mus musculus*), human and cow (*Homo sapiens-Bos taurus*). Additionally, we collect orthologous for seven species with a shared tree (see Figure below). The species are *Homo sapiens, Rhinopithecus bieti, Pan troglodytes, Macaca fascicularis, Mus musculus, Bos taurus, and Sus scrofa.* For the human-mouse dataset we obtained 16,589 protein sequences, while for the human-cow dataset we gathered 15,974 proteins. For the dataset of seven species we collected 11,789 proteins.

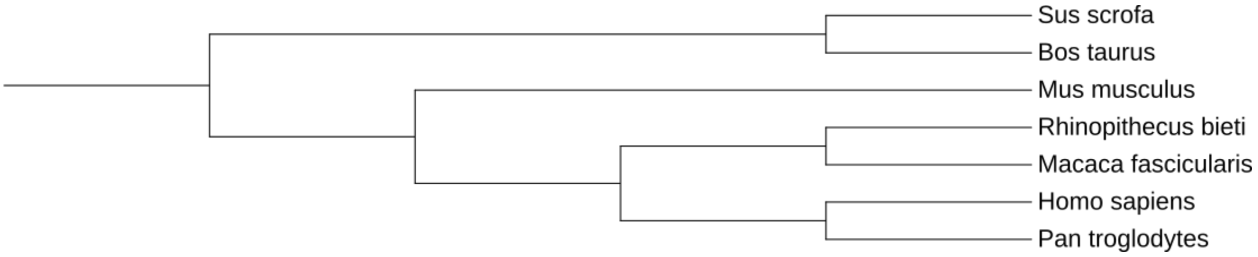


Tree representation of the dataset containing 7 species.

Each set of protein pairs was aligned using the program T-COFFEE (Notredame et al. 2000). Each protein alignment was converted to the DNA alignment using the program pal2nal (Suyama et al. 2006) using the coding sequences for each protein downloaded from the OMA database.

The "amyloids" dataset was obtained by mapping each dataset with the Amypro database (Varadi et al. 2018) to identify proteins with experimental evidence to form amyloids without mutations. The final dataset of 81 amyloidogenic proteins was built by manually adding amyloid proteins curated from the bibliography. The remaining human proteins in each dataset were considered the "reference" dataset.

**Obtaining the evolutionary rates**

Evolutionary rates, expressed as the number of nonsynonymous substitution rates per site (dN), were estimated using the program YN00 and CODEML for the case with seven species with a common tree. Both programs are from the package PAML (Yang 1997).

**Dataset annotation**

The canonical human proteome was annotated with several parameters mapped or obtained from different sources. The different parameters are:

Uniprot ID
Category (Amyloid, Reference / Amyloid, Reference_Cytoplasmic, Reference_Membrane, Reference_Secreted)
Protein Abundance
Protein expression level
Rates measure as d*N* (BH, MH, 7)
APR (*aggregation prone regions*) measured as TANGO
Presence of disulphide bonds
Cell Location
Total protein-protein interactions
Disorder content
Length
Conformational Diversity measured as maximum RMSD100
ΔG0 unfolding
Melting Temperature
Supersaturation (σu (for the unfold state) and σf (for the fold state).

All these variables, precedence and citation can be found in spreadsheet 1.
All the information regarding the amyloid proteins can be found in spreadsheet 2.

**Statistical Analysis and Figures**

All the statistical analyses were performed using R Language and Julia Language.

**Supplementary Figures**

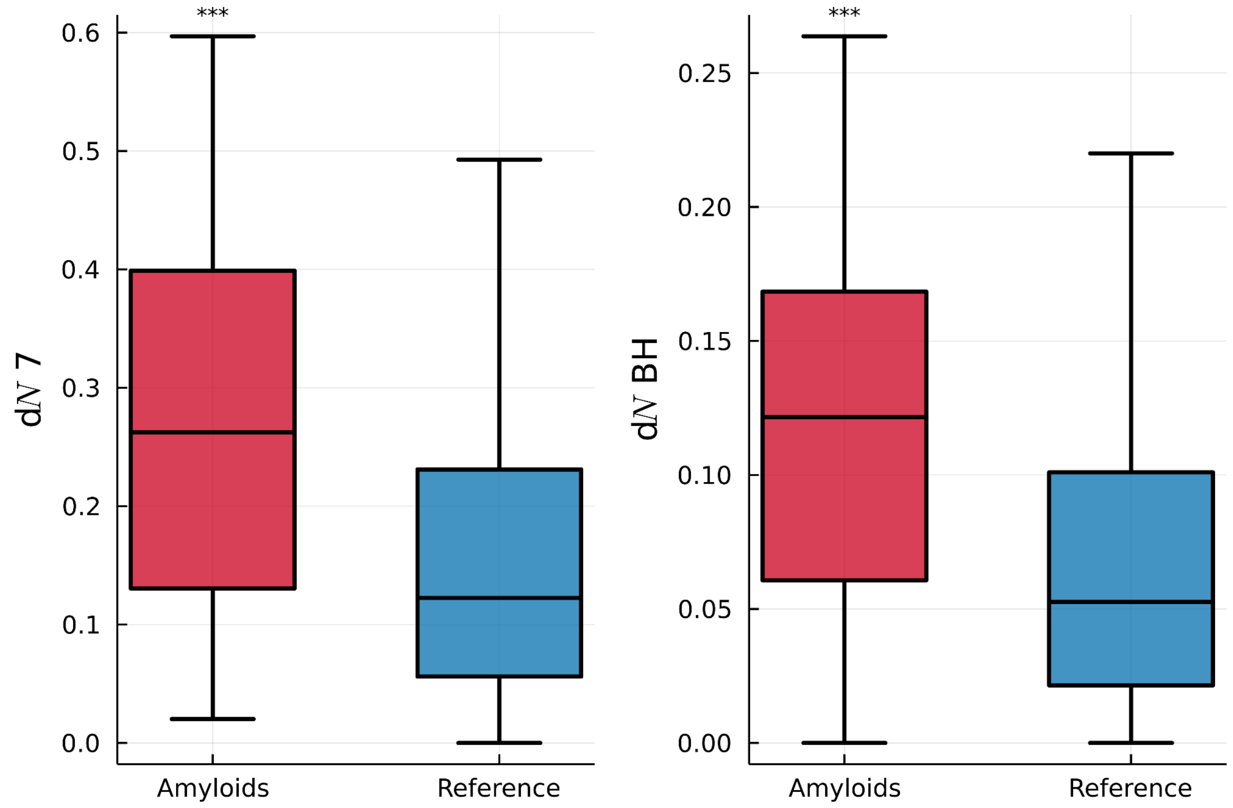


Supplementary Figure 1. Analysis of evolutionary rates with a 7 species dataset (*Homo sapiens, Pan troglodytes, Macaca fascicularis, Rhinopithecus bieti, Bos taurus, Sus scrofa* and *Mus musculus,* right panel, Amyloids n = 43, Reference n = 9,137) and with the *Homo Sapiens - Bos taurus* dataset (left panel, Amyloids n = 61, Reference n = 13,987). Stars indicate Wilcoxon rank-sum test results: ***: p-value<= 0.001; ** p-value <= 0.01; * p-value <= 0.05; ns: p > 0.05.

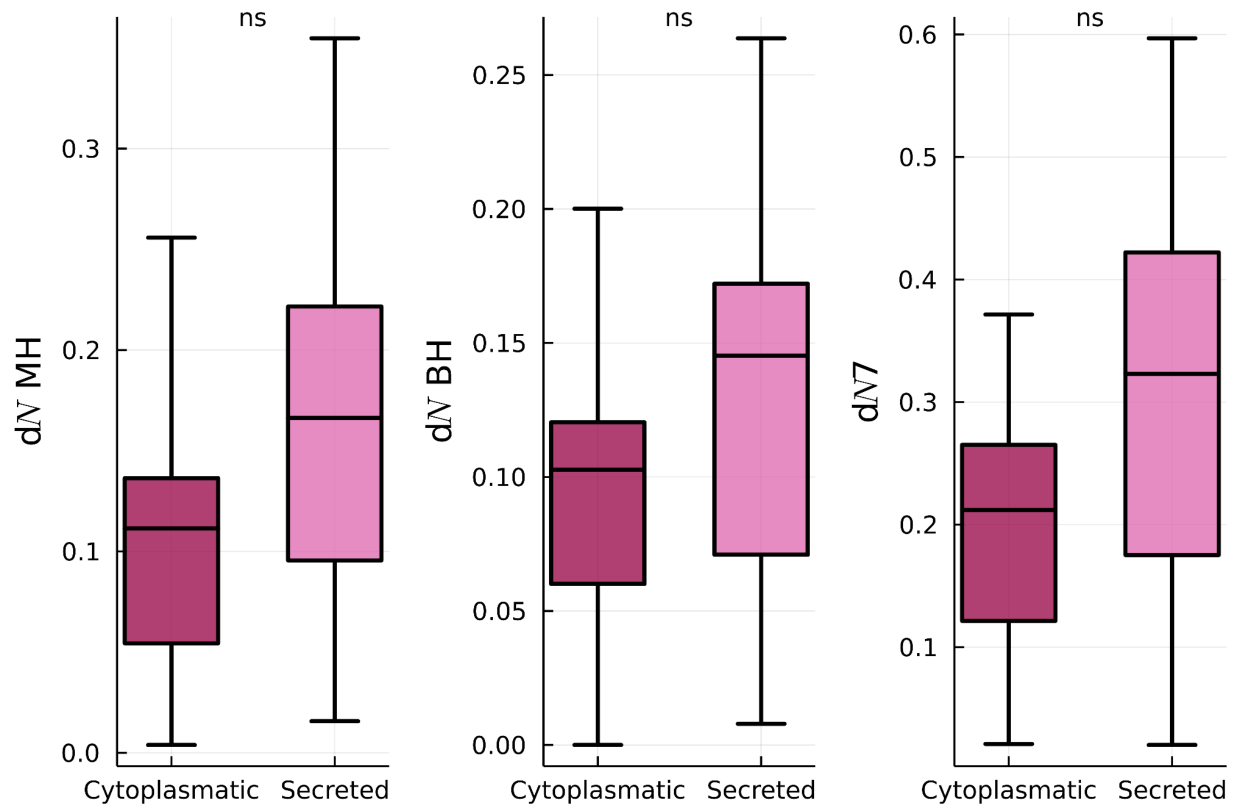


Supplementary Figure 2: Cytoplasmic and Secreted Amyloids show no difference regarding evolutionary levels. Stars indicate Wilcoxon rank-sum test results: ***: p-value<= 0.001; ** p-value <= 0.01; * p-value <= 0.05; ns: p > 0.05.

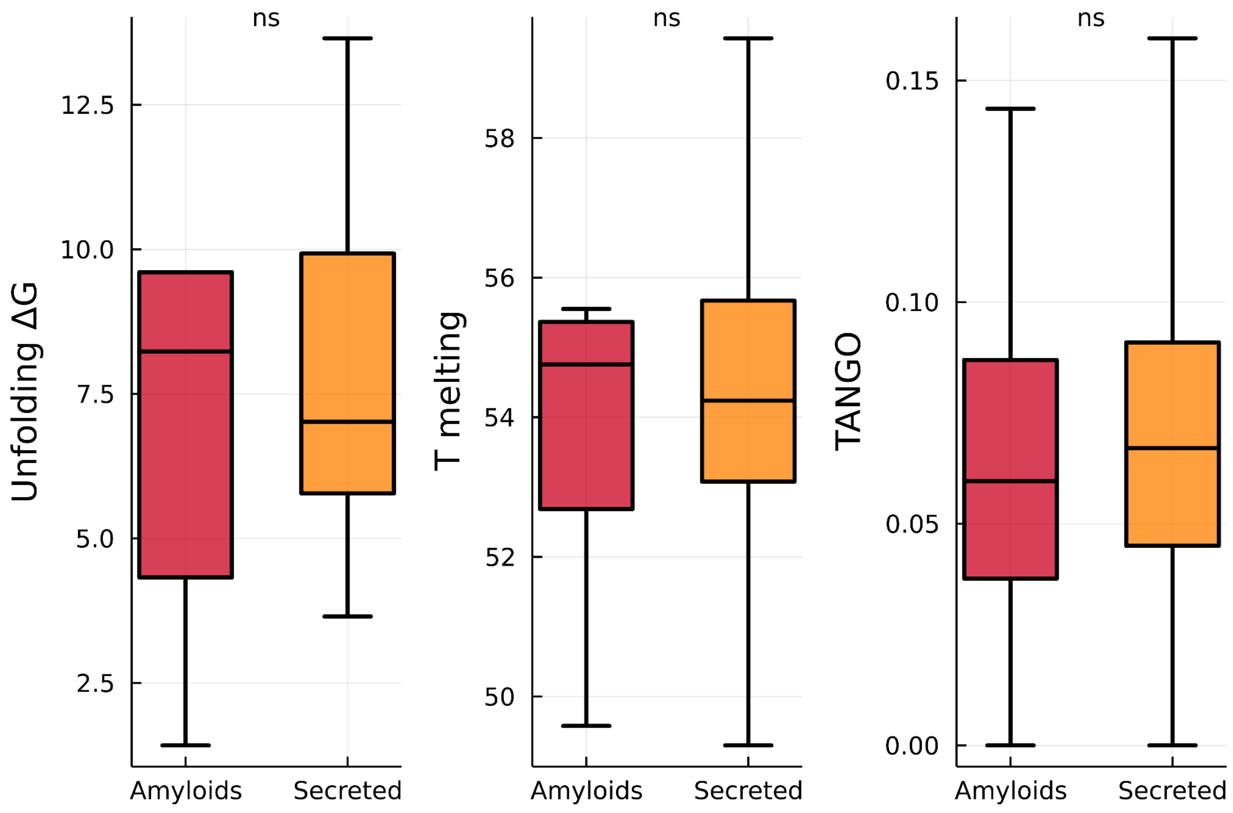


Supplementary Figure 3. Unfolding Gibbs free energy, Temperature melting and TANGO total APR score for both amyloids and secreted proteins. Stars indicate Wilcoxon rank-sum test results: ***: p-value<= 0.001; ** p-value <= 0.01; * p-value <= 0.05; ns: p > 0.05.

**Supplementary Table 1:** Partial correlations results with dN (ρ-p-value) for the amyloid dataset. Parameters with p-values < 0.05 and $R^2$ > 0.2 are shown in bold.
Log(parameter)=log10(parameter+0.001)

| Parameter<br><br>Dataset | Abundance<br><br>ρ<br>p-value | Length<br><br>ρ<br>p-value | RMSD100<br><br>ρ<br>p-value | Protein-protein interactions<br>ρ<br>p-value | Disorder<br><br>ρ<br>p-value | Number of proteins |
|---|---|---|---|---|---|---|
| 7 species | **0.504**<br>**0.0325** | -0.064<br>0.798 | **0.501**<br>**0.0338** | **-0.406**<br>**0.0937** | 0.344<br>0.1611 | 22 |
| Mouse-Human | **0.4703**<br>**0.0314** | -0.161<br>0.484 | **0.493**<br>**0.0228** | **-0.394**<br>**0.0767** | 0.337<br>0.134 | 25 |
| Cattle-Human | 0.377<br>0.100 | -0.1908<br>0.4202 | **0.457**<br>**0.0424** | **-0.495**<br>**0.0261** | 0.2209<br>0.349 | 24 |

**Supplementary table 2:**

Linear Regression models (log(d*N*)~log(RMSD100)+log(Abundance))

| Dataset | Adjusted $R^2$ | p-value |
|---|---|---|
| 7 species | **0.229** | 0.032 |
| Mouse-Human | **0.234** | 0.0206 |
| Cattle-Human | 0.088 | 0.145 |

Linear Regression models (log(d*N*)~log(RMSD100)+log(Abundance)+ log(protein-protein-interactions))

| Dataset | Adjusted $R^2$ | p-value |
|---|---|---|
| 7 species | **0.249** | **0.045** |
| Mouse-Human | **0.2566** | **0.0263** |
| Cattle-Human | **0.3085** | **0.01537** |

**Supplementary table 3:**
Linear Regression models (log(d*N*)~log(RMSD100)+$\sigma_f$)

| Dataset | Adjusted $R^2$ | p-value | n |
|---|---|---|---|
| 7 species | **0.2973** | **0.0477** | 16 |
| Mouse-Human | **0.3769** | **0.0112** | 18 |
| Cattle-human | 0.4346 | 0.0096 | 16 |
